# The Xenopus respiratory system reveals common tetrapod mechanisms for growth, regeneration and healing

**DOI:** 10.1101/2024.10.10.617686

**Authors:** Shiri Kult Perry, Nikko-Ideen Shaidani, Marko E Horb, Neil Shubin

## Abstract

In recent years, we have seen a significant increase in our understanding of the mechanisms of development, regeneration, and healing of the respiratory system. However, most of these studies have been limited by their focus on mammalian systems. Here, we aimed to identify the underlying molecular mechanisms that are active during lung growth and tissue repair in amphibians, specifically Xenopus tropicalis (*X. tropicalis*). First, we analyzed the stem cell composition and signaling pathways that are active in epithelial and mesenchymal cells during lung growth. Then, we established a protocol for lung injury to assess the types of stem cells underlying tissue repair. In mammals, Sftpc+ (AT2) cells are alveolar stem cells that can differentiate to Krt8+ cells during lung homeostasis and post-injury repair. In this study, we identified Sftpc+ cells and Krt8+ cells, along with the activity of key developmental signaling pathways, Hippo and Wnt, during lung maturation at post-metamorphosis stages. We then established a protocol for lung injury using chemically induced injury with bleomycin, which damages the lung through oxidative stress. The results show an elevation in collagen post-injury, indicating bleomycin’s effect in causing lung fibrosis. *X. tropicalis* froglets survived 42 days post-injury, with a continuous decrease in fibrosis. To explore this effect, we analyzed the distribution of lung stem cells; Sox9 protein levels and Sftp gene expression were downregulated at the alveoli 42 days post-injury. The decrease in stem cell marker expression 42 days post-injury suggests they are differentiating as part of the healing process. Nevertheless, we could still detect them after a few weeks of healing. These results suggest that *X. tropicalis* has a regenerative capacity for lung tissue repair and that the same signaling pathways and stem cells are active in both amphibians and mammalian lungs during lung growth, regeneration, and healing. These findings show for the first time the physiological similarities between the anuran and the mammalian lung during growth and tissue repair processes, suggesting *X. tropicalis* as a potential animal model to study lung regeneration.

## Introduction

In recent years we have seen a significant increase in our understanding of the embryonic development and regeneration of the respiratory system; however, most of these studies have focused on mammalian systems (Kotton and Morrisey., 2014, Basil MC, Katzen J, Engler AE *et al*., 2020). Other tetrapods, such as amphibians, also develop a functional respiratory system, which has been studied to a limited extent. Previous work showed that the pathways that control the initial steps of the respiratory system’s development are conserved between mammals, amphibians, and fish (Tatsumi *et al*., 2016, Rankin *et al*.,2015). Despite the importance of these studies, there is still limited knowledge about the later stages of lung development in amphibians, specifically of lung maturation and growth, and the potential for lung regeneration in this animal model. Studying the amphibian respiratory system and its regenerative capacity can lead to significant implications on major respiratory pathologies in humans by elucidating their underlying molecular mechanisms. Here, we aim to identify the signaling pathways that are active during lung growth and regeneration in amphibians, specifically in frogs (anurans).

Xenopus and axolotls have been widely used to study the regeneration of diverse organ systems (e.g., limb and tail, Kragl M, Knapp D, Nacu E, *et al*. 2009, Love, N.R., Chen, Y., Bonev, B. *et al*., 2011, Hayashi S, Kawaguchi A, Uchiyama I, *et al*. 2015), however the mechanisms that regulate the regeneration of some internal tissues, including endodermal tissues such as the lung, remain unknown. Like other organs that originate from the endoderm, such as the liver and pancreas, the lung is a facultative regenerative tissue: following injury, the lung reacts vigorously to replace cells that were lost by re-entering the cell cycle, however normally it is quiescent (Kotton and Morrisey., 2014). This process involves multiple cell types that respond to injury, both in mammals and axolotls (Jensen *et al*., 2021; Kotton and Morrisey, 2014). To date, studies of lung regeneration among amphibians have been scarce. One of those studies has addressed mechanisms of lung regeneration, showing epidermal growth factor receptor family (ERBB) function in lung regeneration in axolotls (Jensen *et al*., 2021). However, despite its importance, it is still unclear if Xenopus lungs can regenerate, and if so, which cell types and signaling pathways are involved in lung regeneration in amphibians, specifically in frogs, such as Xenopus tropicalis (*X. tropicalis)*.

While Xenopus lung embryonic development has been described (Rankin *et al*.,2015), the extent to which these organs recover from injury and renew remains unknown. Specifically, are the signaling pathways that are active during lung growth and maturation conserved between amphibians and other tetrapods, i.e., mammals. By comparing lung developmental and regenerative processes in different animal models, such as amphibians and mammals, enables to study the differences between them and to uncover novel molecular pathways that promote lung renewal. As the lung blood–gas barrier morphology is highly conserved among vertebrates and since it includes the alveolar epithelium that gives rise to lung stem cells in mammals (West, 2011), we focused in this study on these cell types and explored their contribution to regenerative processes in the amphibian Xenopus tropicalis.

Pulmonary fibrosis (PF) is a fibrotic interstitial lung disease (ILDs) that is primarily characterized by inflammation and fibrosis (Cottin *et al*., 2019, Spagnolo *et al*., 2021 Wijsenbeek & Cottin, 2020). During pulmonary fibrosis in humans, there is inflammation and extracellular matrix collagen accumulation. Environmental factors such as air pollution and tobacco smoking, as well as genetic factors, are part of the etiology of PF (Wang *et al.,* 2024). As PF becomes a chronic condition, it progresses to respiratory failure and is, therefore, lethal (Lederer & Martinez, 2018). Although there are treatments to slow down the course of the disease, it remains incurable (Mei *et al*., 2022). Bleomycin (BLM) is a chemical that has been widely used in mammalian animal models, to induce pulmonary fibrosis. In rats and mice, the peak response to BLM lung injury is day 14, with the appearance of fibrosis: First, the elevation of pro-inflammatory cytokines, followed by increased expression of pro-fibrotic markers (fibronectin, procollagen-1), with a peak at ∼day 14. A “switch” between inflammation and fibrosis occurs around day 9 after bleomycin intake (Chaudhary *et al*., 2006, Moeller *et al*., 2008). In humans, bleomycin has been used to treat various malignant cancers, with the possible adverse effect of pulmonary fibrosis, which includes higher expression of fibrogenic factors e.g., (TGF)-beta, connective tissue growth factor, and platelet-derived growth factor (PGDF)-C (Brandt and Gerriets, 2023).

In this work, we outline the development of post metamorphic lungs of Xenopus tropicalis and develop an experimental system to explore lung growth, regeneration and healing processes. Using chemically induced injury of the lungs with bleomycin, known to damage the lung through oxidative stress, we aimed to assess if the amphibian lung regenerates and to identify the cell populations that contribute to organ recovery post-injury.

## Results

### Histology of the Xenopus tropicalis lung at post metamorphic stages

To better understand the morphology of the amphibian lung, and specifically how the lung of Xenopus tropicalis matures, we first performed histological analysis at several stages after metamorphosis using 1-month-old froglets, 4-month-old juveniles and 8-month-old juveniles (Figure 1). We sought to compare an early stage that is closer to metamorphosis, such as a 1-month-old froglet, to later stages that have reached sexual maturity, which in *X. tropicalis* occurs between 4 and 8 months, depending on the gender (Lane *et al*., 2022). We assumed the lungs at those stages will represent the fully matured organ.

**Figure 1.**
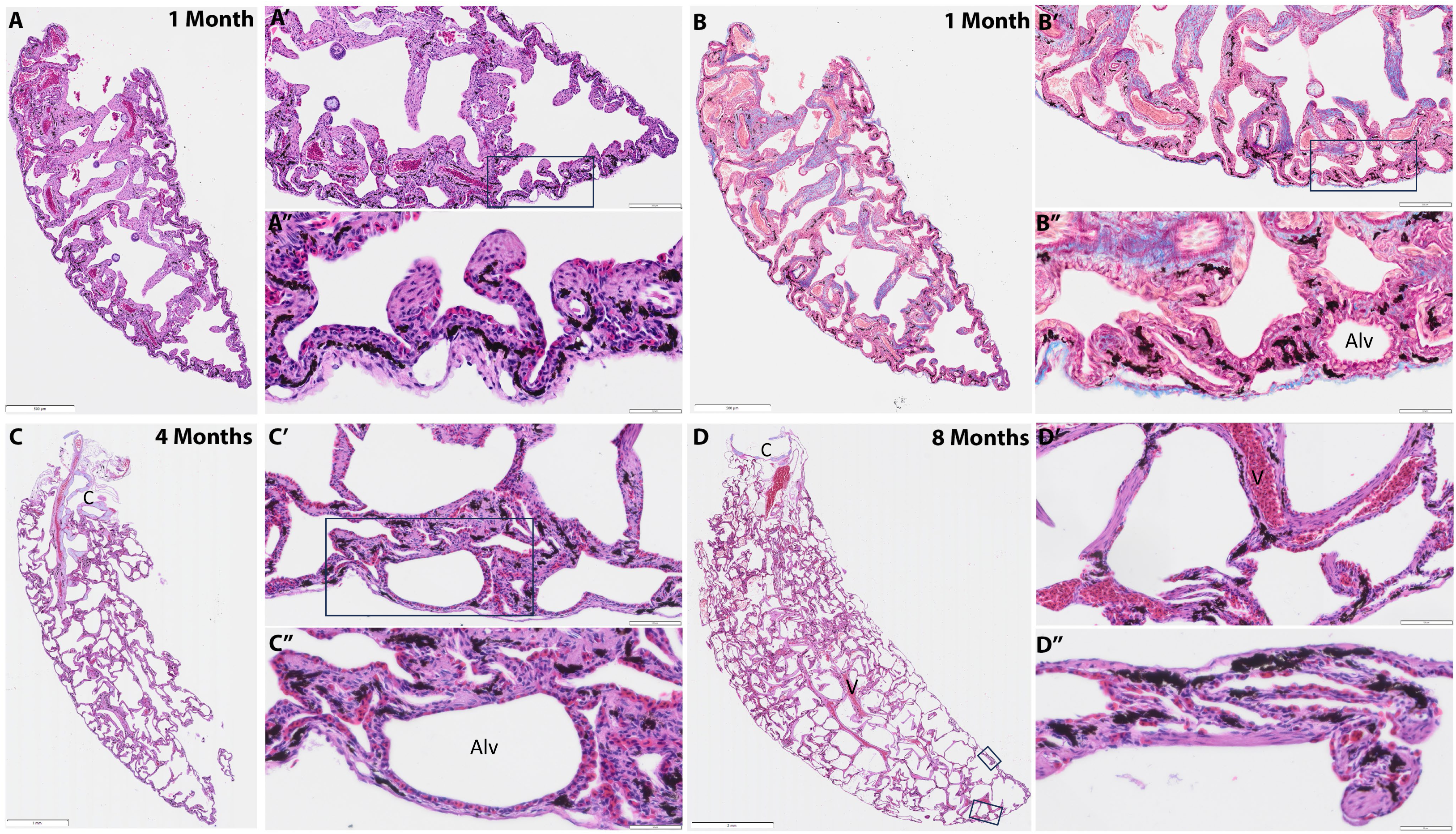
Histology of *Xenopus tropicalis* lungs at post metamorphic stages. H&E staining of the lung of a one-month-old froglet (A) and a parallel section stained with Masson’s trichrome (B). The left panel (A-D) shows lower magnification, where the proximal side is in the upper side, and distal side of the lung is in the bottom of the image. The right panel (A’-D’) shows 20X magnification. A detailed view of the boxed areas is shown as A’’-D’’, with 40X magnification of the distal side of the lung. The alveoli (Alv), blood vessels (V), cartilage (C) are labeled in the figures.

The trachea was identified at the proximal side of the lung, as a cartilaginous tissue, with major blood vessels entering the lung. The alveoli, that are the areas designated for gas exchange, are located on the peripheral sides of the lung, while connective tissues and cartilage nodules are located at the center and proximal sides of the lung (Figure 1). Higher magnification of those areas at the distal side of the lung showed the proximity of connective tissues to the alveoli and the red blood cells (RBC) in nearby capillaries (Figure 1.A’’-D’’). Using Masson’s Trichrome staining we analyzed the collagen composition and localization of connective tissues in the lung in its native state across those stages (Figure 1B). By characterizing *X. tropicalis* lung histology, we identified the alveoli are distributed at the periphery of the tissue, as well as the localization of cartilaginous tissues at the proximal side of the lung, similar to the localization of the trachea in mammals. Masson’s Trichrome staining also emphasized the connective tissues that are located at the central part of the lung in *X. tropicalis* (Figure 1B-B’), similar to *X. laevis* histology (Rankin *et al*., 2015). Overall, the histological analysis emphasizes the elongation of the lung through later stages of development, i.e., at different time points of post-metamorphic stages (Figure 1A-D), as demonstrated by organ growth in size over time, and increase in alveoli number (Figure 1D).

### Stem cell localization and developmental pathways activity in the growing *X. tropicalis* lung

The alveolar cells that secrete SFTPC (surfactant protein C), a surfactant that maintains the stability of the lung by reducing the surface tension of air and fluids, are known as AT2 cells. AT2 cells are alveolar stem cells that can differentiate into AT1 cells during alveolar homeostasis and post-injury repair (Raredon *et al.,* 2019). The two cell types differ in their morphology and function in addition to their relative contribution to healing processes from injury. A recent study identified a transitional stem cell that is Krt8+ that precedes the regeneration of AT1 in reaction to injury (Strunz, M., Simon, L.M., Ansari, M. *et al*., 2020). Sox9 is an essential transcription factor during lung development, as distal epithelial progenitors are identified by high expression of Sox9 (Rawlins *et al.,* 2009, Rockich *et al.,* 2013). The further analysis we performed shows Sftpc, Sox9, and Krt8 are expressed in the lungs of 2-month-old froglet (Figure 2A-C). While Sftpc and Sox9 expression was found in epithelial cells, Krt8 expression was broader, and we could identify it in both epithelial and mesenchymal cells (Figure 2C’). These results indicate that there is a high similarity in cell composition of *X. tropicalis* lung when compared to mammalian lung stem cells, as genes known to be expressed in mammalian alveolar stem cells can be identified in *X. tropicalis* lungs and may contribute to tissue hemostasis and tissue repair post-injury.

**Figure 2.**
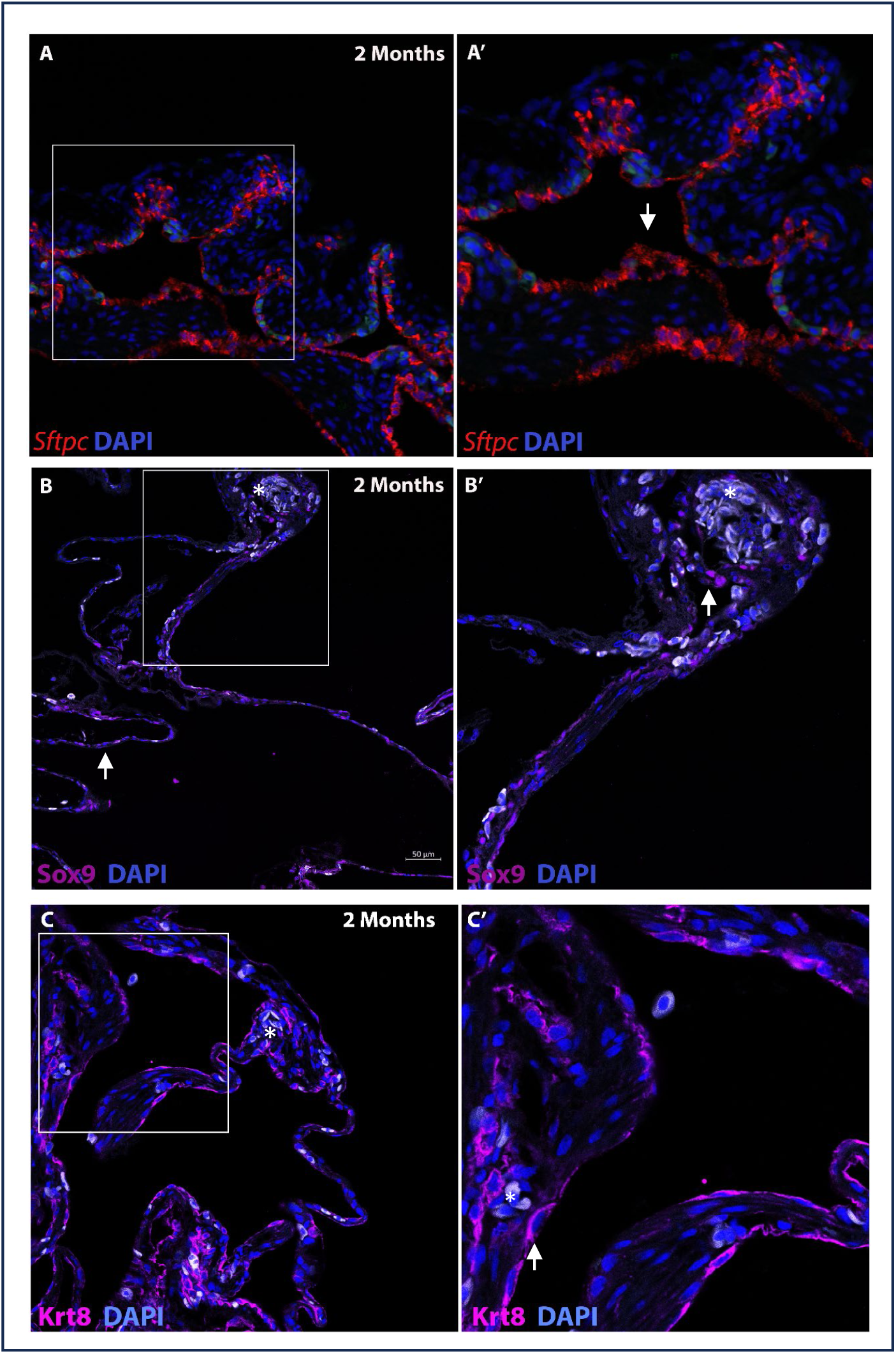
Stem cells localization in the growing *X. tropicalis* lung. (A) HCR *in situ* hybridization of Sftpc (red) and immunofluorescent staining of (B) Sox9 (magenta) and (C) Krt8 (magenta) in 2-months-old froglet. Left panel (A-C) 20X magnification, a detailed view of the boxed areas shown on the right panel (A’-C’). In figures B-C cells shown in white are the red blood cells (examples marked with *).

Previous studies demonstrated the activity of Wnt signaling pathway during endodermal tissue development, including the respiratory system (Rankin *et al*., 2015). Beta-catenin activity is essential for NKX2.1 expression, which leads to the formation of the respiratory system (Kim *et al*., 2019). In mammals, epithelial-mesenchymal crosstalk during lung development is mediated by Wnt (Volckaert & De Langhe, 2015). This pathway is central for stem cell differentiation, such as Sox9+ epithelial cells, and therefore contributes to the tissue regenerative capacity and healing processes. Since both Wnt and Hippo pathways are key developmental pathways that are active in the embryonic and adult mammalian lung (Lange *et al*., 2014., Ostrin *et al*., 2018), we sought to analyze the localization of the cells that are positive to beta-catenin and Yap1, respectively, during the growth and maturation stages of the amphibian lung (Figure 3). At one-month-old froglet stage, high levels of beta-catenin were identified in epithelial cells at the distal side of the lung (Figure 3A). At later stage of 8-month-old froglet, only a few cells expressed beta-catenin, when compared to the earlier stage. At 4-month-old froglet, both pathways, Hippo and Wnt, were found to be active at the alveoli located at the distal side of the lung (Figure 3B). At the earlier stage of 2-month-old froglet, Yap1+ cells were identified at the distal side of the lung (Figure 3CF). Overall, in both early and late stages, cell populations with low or high expression levels of Yap1 were identified in proximity.

**Figure 3.**
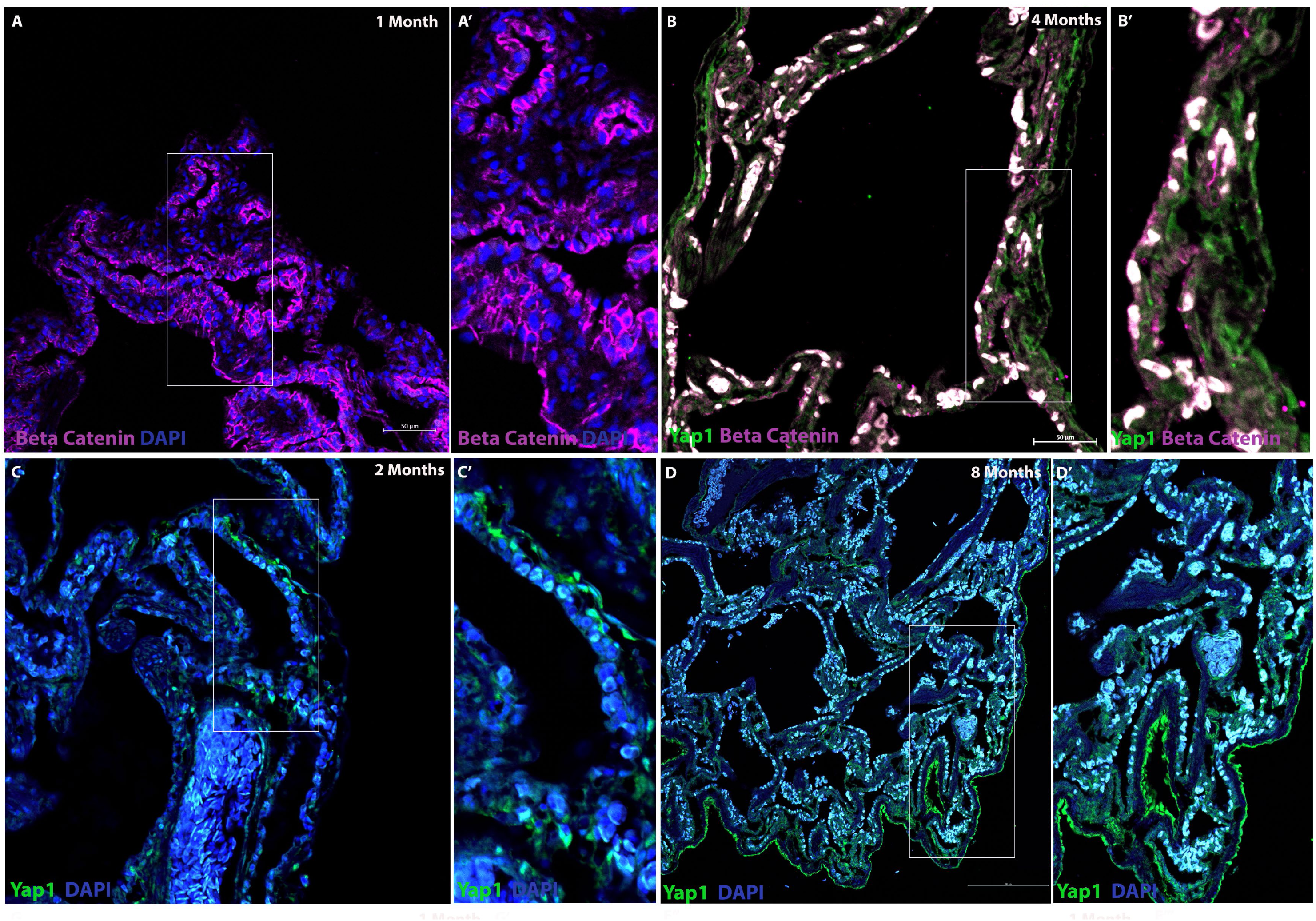
Developmental pathways activity in the growing *X. tropicalis* lung. (A) β-catenin (magenta) in 1-month-old froglet’s lung. (B) β-catenin (magenta) and Yap1 (green) at 4-month-old juvenile lung demonstrating the activity of the two pathways, Hippo and Wnt in adjacent cells. Immunofluorescent staining of; (C) Yap1 (green) in 2-month-old froglet lung and 8-month-old juvenile lung (D). Left panel (A-D) 20X magnification, a detailed view of the boxed areas shown on the right panel (A’-D’). In all figures cells shown in white are the red blood cells (examples marked with *).

Finally, as the canonical Wnt/beta-catenin signaling pathway is known to upregulate Sox9+ expression and promote the maintenance of epithelial stem cells at the distal side of the mammalian lung (Hu *et al*., 2020), we verified that both beta-catenin and Sox9 are active in adjacent cells at the distal side of the amphibian lung in one-month-old froglet (Figure S2). These results suggest that the signaling pathways that promote lung development and maturation are well conserved across amphibians as frogs, and mammals. Additionally, these results indicate the localization of stem cells in the froglet lung at post-metamorphic stages, which may have a role during lung homeostasis and post-injury repair. These results raise the question of the regenerative capacity of the amphibian respiratory system and how it would respond to chemically induced injury, which has been widely used with mammalian lungs.

### The amphibian lung shows an increase in ECM deposition post injury that reduces during healing

To test the regenerative capacity of the amphibian respiratory system, we used bleomycin to induce lung injury in *X. tropicalis* chemically and followed the healing process of lungs post-injury in froglets. Late-stage *X. tropicalis* tadpoles (St. 65-66) were grown, underwent metamorphosis to froglets, and were treated with bleomycin as the injury protocol. Xenopus is known to retain its regenerative capacity for early life stages in various tissues and then lose this capability for some tissue types after metamorphosis. Therefore, we chose a stage that we hypothesized to be optimal: with lungs but close to metamorphic stages. We collected lungs from control and bleomycin-treated froglets at 4 time points: 7,14, 21-, and 42-days post-injury (dpi). Masson’s trichrome staining revealed an elevation in collagen deposition and alveolar enlargement post-injury, indicating bleomycin’s effect in causing lung fibrosis (Figure 4). At 7 dpi, there was no apparent difference in the histology of control and bleomycin-treated lungs (Fig. 4A,B). At 14 dpi, an increase in extracellular matrix (ECM) in the lung after bleomycin treatment (BLM) was observed (Fig. 4C,D, higher magnification shown in C’’-D’’, area marked with arrow). A closer look at 21 dpi show cells accumulating in the lung that underwent BLM treatment (Fig. F’) At a later timepoint of 42 dpi, the ECM seems to be near normal levels (Fig.4G-H, G’-H’, G’’-H’’). Overall, the Xenopus froglets survived 42 days post-injury, with a decrease in fibrosis. To summarize, the findings demonstrated an increase in collagen levels 14 days after injury, highlighting bleomycin’s role in inducing lung fibrosis in the Xenopus model. This was followed by a steady decline in fibrosis, with Xenopus froglets surviving 42 days post-injury. We then aimed to identify the stem cell populations contributing to organ recovery post-injury.

**Figure 4.**
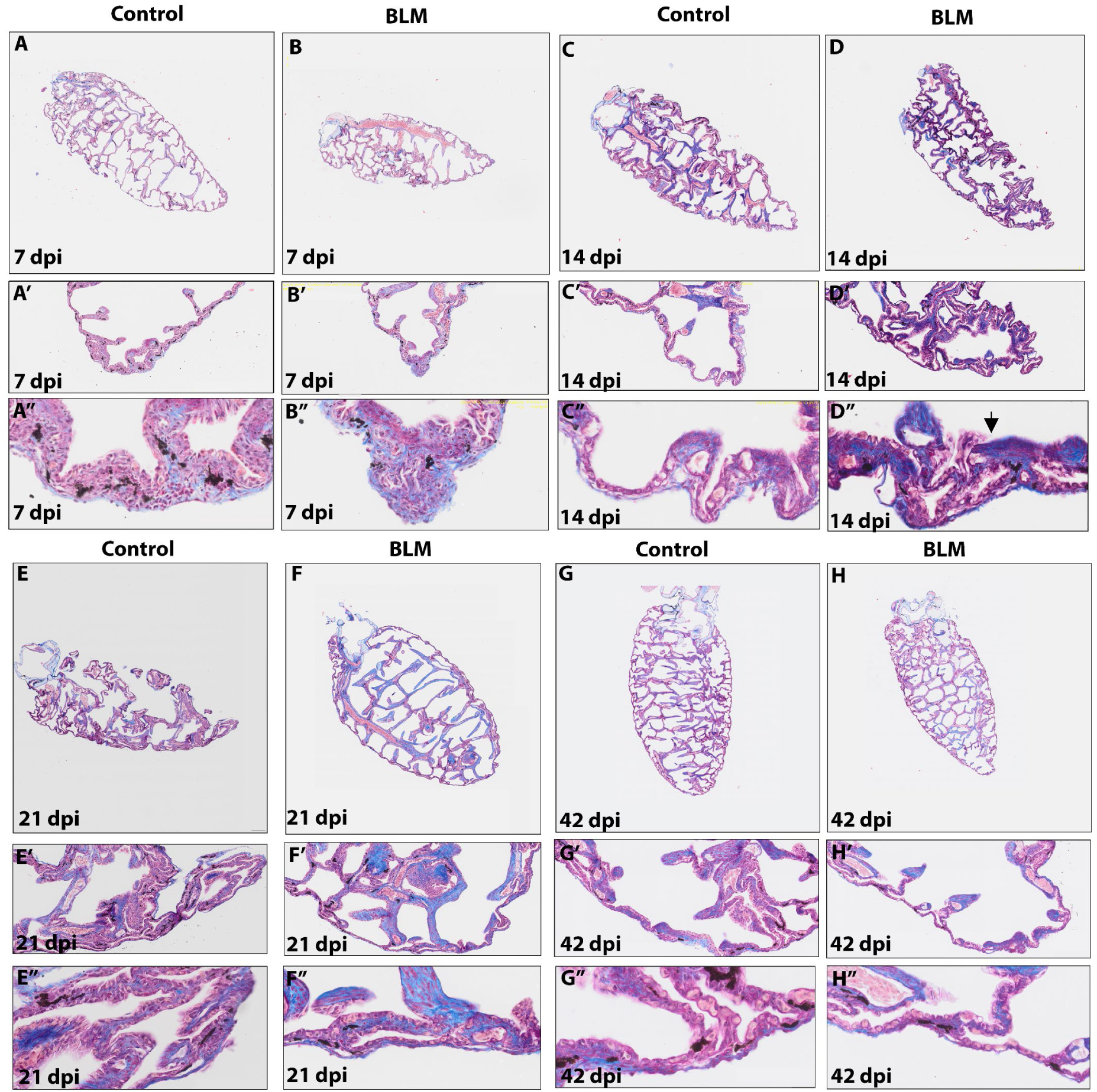
Masson’s Trichrome analysis of Xenopus tropicalis lungs post injury. (A-H) The lungs of PBS injected animals (Control) and Bleomycin injected animals (BLM) were analyzed 7,14,21 and 42 days post injury (dpi). Masson’s trichrome staining of froglet’s lungs show collagen fibers in blue, suggesting there is an increase in extra cellular matrix (ECM) in the lung after bleomycin treatment (BLM). For each timepoint, top to bottom: 1.6X magnification (A-H), 10X magnification (A’-H’), 40X magnification (A’’-H’’) of the distal side of the lung. Arrow indicate the areas where there is an increase in ECM (blue) at 14 dpi (D’’).

### Stem cells and ECM distribution during healing from injury

The protocol we established to chemically induce lung injury using bleomycin with amphibians enabled us to assess lung regenerative capacity in Xenopus tropicalis. To explore the healing process, we analyzed the distribution of lung stem cells, as well as the distribution of Collagen I expressing cells (Figure 5). We compared Sox9 protein levels in control vs. BLM treatment at 42 dpi. In parallel, we compared the distribution of Collagen I expressing cells. Sox9+ cell numbers were lower in BLM-treated lungs than in the control. Collagen I expressing cells, however, were distributed in a similar manner in both conditions (Figure 5B, B’), in agreement with the Masson’s analysis we performed (Figure 4), suggesting that by 42 dpi, there is a reduction in fibrosis. Next, we performed HCR *in situ* hybridization for Sftp gene expression (Sftpc, Sftpb) that are known to be expressed in lung alveolar epithelial stem cells (AT2). The results show that Sftpc and Sftpb mRNA expression is modified at the alveoli during healing (Figure 5D-E, D’-E’). While the Sftpb mRNA expression is clearly reduced at 42 dpi, Sftpc mRNA expression is reduced to a lesser extent. The decrease in stem cells marker expression 42 days post-injury suggests they are differentiating as part of the healing process. Nevertheless, we could still detect them after a few weeks of healing. Hence, these results suggest that the abundance of alveolar stem cells in a relatively small tissue is the mechanism underlying the recovery of *X. tropicalis* after lung injury.

**Figure 5.**
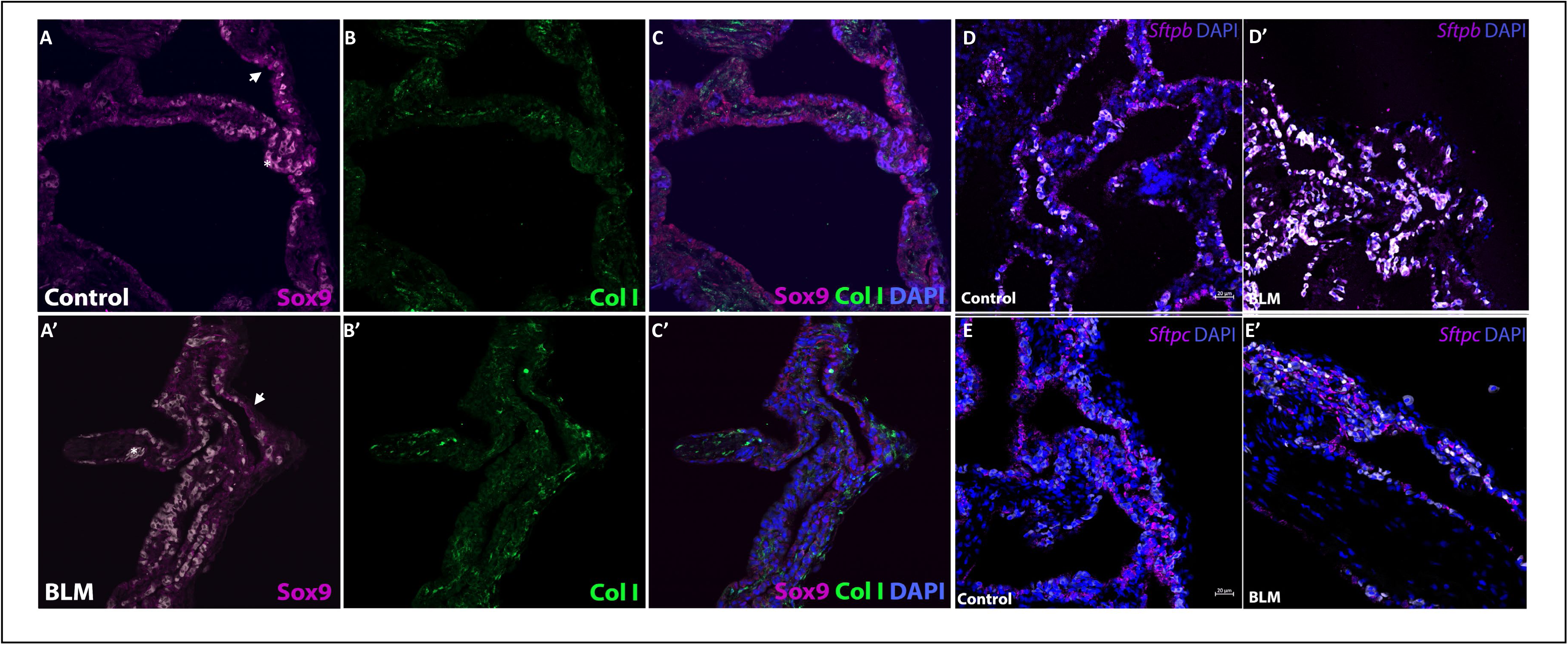
Stem cells distribution during healing from injury. (A) Immunofluorescent staining of control and bleomycin treated froglet’s lung (BLM) for (A) Sox9 (magenta), and (B) Collagen I expressing cells (green) and (C) their merge images demonstrating the recovery of the tissue as stem cells identified in the alveoli, and connective tissue localized in the periphery of the tissue, i.e., in adjacent cells at 42 dpi (day post injury). Sftp gene expression is affected during healing from bleomycin injury: HCR in situ hybridization of (D) Sftpb (magenta) and Sftpc (E, magenta) at 42 dpi in control (Con) and injured lungs (BLM), demonstrating their decrease during healing from lung injury. (A-E, A’-E’) 20X magnification. Cells shown in white are red blood cells (examples marked with *).

## Discussion

Our results suggest that Xenopus tropicalis has a regenerative capacity for lung tissue repair. Overall, we present in this study the physiological similarities between the amphibian and the mammalian lung during growth and tissue repair processes, suggesting Xenopus tropicalis as a potential animal model to study these processes further. To establish a timeline for lung growth in amphibians and follow the cellular processes that lead to its full maturation, we compared the lungs of Xenopus tropicalis in different stages. We searched for various types of lung cell populations and how they differ at the lung at different stages. Ultimately, this analysis allowed us to identify various cell populations that build the growing amphibian lung.

There are key processes that occur in different tissue types following injury. Previous studies demonstrated the key phases of regeneration of some tissues, such as limbs, as wound healing, formation of blastema, and re-development of blastema (Suzuki *et al*., 2007, Stocum 2017). However, regeneration and healing processes are not always binary in their representation, as in the case of epithelial tissues such as the gut or the lung, where the process seems more gradual and appears differently than a blastema. Unlike mammalian limbs, epithelial tissues of mammals can regenerate to some extent (Iismaa *et al*., 2018, Eenjes *et al*., 2022). Still, the efficiency of these processes may be lower in pathological conditions, such as lung diseases. In this study, we show that amphibians, specifically *X. tropicalis,* have lung regenerative and healing capacities for the first time.

Previous studies investigated the respiratory system of various models to address how the organism developed adaptations of the repertory system according to its environment. Some amphibians, like *X. tropicalis,* undergo metamorphosis and are challenged as they need to breathe and perform gas exchange in aquatic and terrestrial environments. Some amphibians, like the lungless salamanders, have lost their lungs but developed their ability to respire through other organs, such as the buccopharyngeal cavity (Lewis, Dorantes, and Hanken 2018). Therefore, amphibians are a fascinating example of the diversity of physiological solutions and how developmental processes shape them. Another study describes the cell types that reside in the land-dwelling frog (*Microhyla fissipes*), a frog that shifts from an aquatic to a terrestrial environment (Chang, L., Chen, Q., Wang, B. *et al.,* 2024). Among their findings, there is a cross-species comparison of scRNA-Seq data, showing this frog has lung cell types that were not found in other frog species (*X.laevis*), as well as cell types that are similar to mammalian lungs. Altogether, these findings in addition to our study suggest that the study of amphibian lungs and respiration processes can lead to novel findings.

Lung regeneration may aid *X. tropicalis* in its natural environment for several reasons. First, it allows the individual to recover from injury by predators or other individuals from the same species. Second, regeneration may allow recovery from parasites that invade the lungs, such as helminths (worms) (Nichols 2000, Gendron, 2013., Imasuen and Aisien, 2015). In its natural habitat, *X. tropicalis* can host at least ten worm parasites (Imasuen and Aisien, 2015). Digenean is a parasitic flatworm. While the larval form often encysts in the host tissue, the adult is located in the lumen of many host organs, including the lungs (Gendron, 2013). Another possible parasite is adult trematodes, which were reported to be found in the lungs of frogs, but it is unclear if they cause a disease (Nichols 2000). The genus Rhabdias of nematodes is also found in the lungs of amphibians, including frogs (therefore termed lungworms), causing pulmonary damage and additional infections. The adult lives in the lungs and grows using blood and respiratory secretions. In some cases, the larvae can mature on the soil to reach an infective stage. Interestingly, this parasite penetrates the animal’s skin and migrates until it enters the lungs (Nichols 2000). Third, environmental factors, such as pollutants (Chen *et al*., 2022). These are all possible explanations for the evolutionary pressure to develop regenerative organs in amphibians, specifically a regenerative respiratory system.

As the lung blood–gas barrier morphology is highly conserved among vertebrates and since it includes the alveolar epithelium that gives rise to lung stem cells in mammals (West, 2011), we focused in this study on these cell types to explore their contribution to regenerative processes in the amphibian Xenopus tropicalis. However, there are also differences between amphibians and mammals, such as the gross structure of the lung, which is different due to branching morphogenesis in mammals that does not occur in amphibians. The diversity of lung morphologies, cell type diversity, extracellular matrix (ECM) composition, metabolic processes and other physiological differences such as body size, endocrine activity, in addition to the organism’s ecological niche, which includes environmental factors (Ref), are all possible differences which require further investigation on how they affect amphibians ‘respiratory system development and regeneration, and specifically in *X. tropicalis.* By comparing lung regeneration processes in different animal models, such as amphibians to mammals, it is possible to study these differences and to uncover novel molecular pathways that promote lung renewal. Our results demonstrate the induction of lung injury in *X. tropicalis* using bleomycin to study tissue repair. The results suggest that developmental processes that are active during lung growth are evolutionarily conserved between amphibians and mammals, hence the possibility of further exploring the Xenopus lung as a disease model for human respiratory pathologies.

Studying the amphibian respiratory system can have significant implications on major respiratory pathologies in humans by elucidating their underlying molecular mechanisms. Previous studies on limb regeneration in amphibians demonstrated how immune cell activity differs from humans by promoting tissue renewal (Godwin *et al*., 2013). Therefore, studying immune responses during lung injury in amphibians makes it possible to identify the underlying regulatory mechanisms and develop new therapeutical treatments. Altogether, this strategy can uncover novel therapies for human lung disease that involve inflammation, such as pulmonary fibrosis (PF), Chronic obstructive pulmonary disease (COPD), acute respiratory distress syndrome (ARDS) that is related to Covid-19, lung transplantation and cancer, as well as chronic lower respiratory diseases (Burney *et al*., 2017, Torres Acosta & Singer, 2020, Wijsenbeek & Cottin, 2020). Finally, by comparing our findings to previous studies on mammalian lung development and regeneration, we can reveal common mechanisms that have evolved in nature to allow lung regeneration. Thus, our study sheds light on this fascinating yet poorly understood aspect of respiratory system regeneration.

To conclude, this study links cell biology, evolution, and developmental biology to uncover the molecular machinery that is active during lung growth and regeneration in amphibians, specifically Xenopus tropicalis. This study paves the way to new applications of this animal model, which can lead to new strategies for studying lung development, physiology and evolution.

## Materials & Methods

### Animals and Ethics statement

Animals were purchased from the National Xenopus Resource (NXR, Marine Biological Laboratory RRID:SCR_013731) or Xenopus1. The study involved *X. tropicalis* (NXR_1018) NF57 tadpoles to juvenile frogs staged using Nieuwkoop and Faber (NF) nomenclature (www.xenbase.org RRID:SCR_003280, Fisher *et al*. 2023). Animals were housed in a temperature-controlled facility with 12:12-h light: dark cycles, using recirculating water system or stationary water tanks, and a commercial diet (Sera micron powder or pellets, depending on animal stage). In all experiments, *n*≥2 froglets or juvenile frogs were used.

Ethics Animal experimentation: All animals were maintained and used in accordance with protocols approved by the University of Chicago Institutional Animal Care and Use Committee (IACUC; permission no. 72696). All animals used in this study were housed in an Animal Care Facility at the University of Chicago. Animals were monitored daily for food and water intake and were inspected weekly by the head of the animal facility in consultation with the veterinary staff.

### Chemical induction of lung injury

Animals were anesthetized in buffered 0.03-0.05% tricaine methanesulfonate (MS222) until the righting reflex was lost. Animals were transferred to a moist gauze-covered plate. Bleomycin (0.01-0.025 units) was then injected intraperitoneally to induce lung injury in *X. tropicalis*. The control group of animals was administered with PBS injected intraperitoneally. Animals were monitored for recovery and placed back into their housing, that included a platform for them to rest close to water surface while they recover. After injury with bleomycin, animals were inspected daily and euthanized at specific time points with 0.5% MS-222.

### Histology, Masson’s Trichrome staining & Cryo-sectioning

Harvested lungs were fixed in 4% paraformaldehyde (PFA)/PBS overnight at 4C with constant agitation, then washed gradually with ethanol to 100% for dehydration and embedded in paraffin. Tissues were sectioned at a thickness of 5 µm, then stained with H&E (hematoxylin/eosin) or Masson’s Trichrome. Slides were imaged using Olympus VS200 Slideview Research Slide Scanner.

### HCR RNA-FISH

HCR protocol was performed using paraffin sections as described previously (FFPE tissue sections, Choi *et al*., 2018). Single-and double-fluorescent HCR *in situ* hybridization on paraffin sections was performed using 546- and/or 647-fluorescently labeled probes (listed in Table 1). Slides were mounted using Fluoromount-G Mounting Medium, with DAPI (Invitrogen, 00-4959-52). Slides were imaged using Zeiss LSM 710 or LSM 900 confocal microscope.

### Immunofluorescence

Paraffin sections (5 µm thickness) were deparaffinized and rehydrated gradually, washed briefly in PBSTr (PBS+0.1% Triton X-100) then underwent antigen retrieval with EDTA pH=9 solution that was boiled in a microwave, and cooled down to room temperature. After two washes in PBSTr, to reduce background fluorescence, sections were then incubated with pre-cooled sodium borohydride (1 mg/ml in PBS), for three consecutive 10 min incubations without any intermediary washes. The sections were washed twice briefly with PBSTr. For blocking, sections were incubated in 10% horse, 1% BSA serum dissolved in PBT (PBS+0.1% Tween20) solution for 1 hour in room temperature. The sections were then incubated overnight in 4C with the primary antibody solution as follows: Sox9 (1:150, AB5535, Millipore Sigma), β-catenin (1:200, 17565-1-AP, 66379-1-Ig, Proteintech), Yap1 (1:200, 13584-1-AP, Proteintech), Krt8 (1:200, 27105-1-AP, Proteintech),), Collagen type I (SP1.D8 DSHB, 19 ug/mL). The sections were washed with PBSTr and incubated with secondary fluorescent antibodies (1:200, 715-165-150, 715-175-151, 711-166-152, JacksonImmunoResearch or 1:200, 647 anti-rabbit, Invitrogen) for 1 hour at room temperature or 4C overnight. For double fluorescent immunostaining, the sections were washed three times in PBSTr and incubated overnight in 4C with the primary antibody solution. The sections were washed with PBSTr and incubated with secondary fluorescent antibodies for 1 hour at room temperature. Slides were mounted using Fluoromount-G Mounting Medium, with DAPI (Invitrogen, 00-4959-52). Slides were imaged using Zeiss LSM 710 or LSM 900 confocal microscope.

## Acknowledgments

The authors thank all past and present members of the Shubin laboratory for advice and support. We thank Emily Hillan and Yulia Schwartz for discussions and comments on the draft of the manuscript. We thank Dylan Jockel, Melvin Bonilla, Emma Heras and Ophelia Dominguez for animal care support. The authors thank the University of Chicago Animal Resources Center (RRID:SCR_021806), the Culver animal house facility staff, Dr. Darya Mailhiot, Dr. Faazal Rehman, Juan Rodriguez, and Courtney Losser for their assistance with animal care and housing. The authors thank the University of Chicago Human Tissue Resource Center (RRID:SCR_019199), and the University of Chicago Integrated Light Microscopy Core (RRID: SCR_019197). We thank Marko E Horb, Nikko-Ideen Shaidaini, and James Parente (National Xenopus Resource [NXR], Marine Biological Laboratory) for husbandry and providing *X. tropicalis* tadpoles and to Robert Weymouth from Xenopus1. The SP1.D8 developed by Furthmayr, H., Yale University, was obtained from the Developmental Studies Hybridoma Bank, created by the NICHD of the NIH and maintained at The University of Iowa, Department of Biology, Iowa City, IA 52242. This work was supported by the University of Chicago Biological Sciences and the Brinson Foundation (N.S); and by grants from NIH P40OD010997 and R24OD030008 (M.E.H); and by the University of Chicago, the Chicago fellows program (S.K.P). This work was supported in part by the Zuckerman STEM Leadership Program (S.K.P).

